# Cleaved vs. Uncleaved: How Furin Cleavage Reshapes the Conformational Landscape of SARS-CoV-2 Spike

**DOI:** 10.1101/2025.03.12.642945

**Authors:** Natesan Mani, Raghavendran Suresh, Srirupa Chakraborty

## Abstract

The SARS-CoV-2 Spike protein is the primary target for vaccine design, with immunogens typically engineered to enhance stability by introducing proline mutations (2P) and mutating or deleting the Furin Cleavage Site (FCS). While these modifications improve structural integrity, studies suggest that furin cleavage can play a functional role in Spike protein dynamics, potentially enhancing ACE2 receptor binding. However, the impact of this cleavage on the unbound form of the Spike protein remains unclear. In this study, we use extensive all-atom molecular dynamics (MD) simulations to compare the structural and dynamic properties of cleaved and uncleaved Spike proteins in their pre-fusion, unbound state. Our results show that Furin cleavage significantly alters allosteric communication within the protein, increasing correlated motions between the Receptor Binding Domain (RBD) and N-terminal Domain (NTD), which may facilitate receptor engagement. Principal Component Analysis (PCA) reveals that the cleaved and uncleaved Spike proteins sample distinct conformational landscapes, with the cleaved form displaying enhanced flexibility and a broader range of RBD tilt angles. Additionally, Furin cleavage primes the S2 subunit by expanding the central helix, potentially influencing the transition to the post-fusion state. Glycan clustering patterns further suggest an adaptive structural response to cleavage, particularly in the NTD and RBD regions. These findings highlight the potential functional consequences of FCS deletion in immunogen design and underscore the importance of considering the native cleavage state in vaccine and therapeutic development.

## Introduction

The Covid-19 pandemic, caused by the SARS CoV-2 virus, affected around 700 million people around the world and led to a loss of almost 6 million lives around the world since 2019(1). The first case was reported in Wuhan and from then on, there have been countless mutations of the virus. While most of these mutations have had little or no impact, a few have increased the virus’s ability to infect and spread, higher tolerance to vaccines and increased severity. These have been labelled as the Variants of Concern (VOC). So far, the notable VOCs have been Alpha, Beta, Delta, and Omicron[7]. All these variants have a higher rate of transmissibility than the original strain, with the Delta variant causing the most deaths among all other variants for people who were not vaccinated. The Delta variant also infected those vaccinated, leading to booster shots being administered.

The Spike protein helps in virial attachment by binding to the receptor on the host cell and also induces membrane fusion(2). It elicits a large majority of responses from a wide range of antibodies(3). This makes it the optimum target for vaccine development and immunogenic design. The key difference between SARS CoV-2 and its predecessor SARS CoV is the presence of a Furin cleavage site (FCS) on the Spike protein(4). The complete FCS consists of a 20 amino acid sequence consisting of the range 672-691. The core region is of the sequence P_681_RRAR_685_, with RRAR constituting the minimal polybasic site required for cleavage. The presence of this cleavage site is one of the major reasons that led to the increased virulence, pathogenesis and transmissibility of the virus as Furin is ubiquitously expressed in humans. Given past success with viral research, the Extra Cellular Domain (ECD) was used for vaccine design and immunogenic studies. However, further stabilization of the ECD to make it soluble and crystallizable. This is performed using two methods: the residues at K986 and V987 of the trimer were replaced by Prolines (2P) to stabilize the trimer and the Furin cleavage site (FCS) is mutated to GSAS or deleted, which helps to keep the Spike protein intact(5-7). However, several studies have also reported that Furin cleavage could confer useful properties to the Spike protein. This occurs because of the destabilizing of the Spike protein post-cleavage, which enhances the ability of ACE2 binding(8-12). Hence it is essential to understand the impact of the Furin cleavage on the overall protein allostery

While the pandemic triggered a tumultuous amount of literature on the structural studies of the Spike protein, structural modifications and the essential dynamics observed because of Furin cleavage have not yet been studied. This is because it is extremely resource-intensive to only use experimental techniques and, in such cases, computational methods such as Atomistic Molecular Dynamics (MD) can help us understand the structure and dynamics. MD can aid experimentalists by providing a quantitative framework that allows us to capture the impact of Furin cleavage on the Spike protein and help us design effective therapeutic strategies.

The Receptor Binding Domain (RBD) consists of residue ranges 319-541 and forms the first point of contact with the ACE2 receptor. It has also been proven that among all the subunits of the Spike protein, the majority of antibodies produced by the host target the RBD. The RBD transitions from an ACE2 inaccessible RBD-Down conformation which is energetically favored, to an ACE2 accessible RBD-Up conformation which is a necessary condition for ACE2 binding.

The Receptor Binding Motif (RBM) on the RBD, is the first point of contact with the ACE2 receptor. In the RBD-Down state, the RBM is buried and hence the interaction of the RBM with ACE2 depends on the transition of the RBD from the Down to Up conformation[9].

To investigate the role of the RRAR motif in shaping Spike protein dynamics, we simulated both RBD-Up and RBD-Down conformations in cleaved and uncleaved states, each for five independent trajectories of ∼1.2 µs. Our analysis revealed that Furin cleavage enhances correlated motions across critical domains, particularly between the Receptor Binding Domain (RBD) and the N-terminal Domain (NTD), suggesting a more coordinated structural response that could influence receptor binding and immune evasion. Principal Component Analysis (PCA) further demonstrated that cleaved and uncleaved systems explore distinct conformational landscapes, indicating that cleavage modulates large-scale structural transitions. Notably, in the RBD-Up conformation, we identified two dominant and stable motion states: a “blocked” state, where the RBD tilts inward, potentially reducing ACE2 accessibility, and an “exposed” state, where the RBD tilts outward, enhancing receptor engagement. The presence of these two stable states in the cleaved conformation suggests that Furin cleavage may fine-tune the balance between antibody evasion and efficient host cell entry, providing critical insights for vaccine and therapeutic design.

## Results

The MD simulations done in this study were performed on the post-cleavage (Cleaved) and pre-cleavage (Uncleaved) conformations of the Spike protein of SARS CoV-2. For each of these systems, two different conformations of the Spike: RBD accessible (RBD Up) and RBD inaccessible (RBD Down) were selected. The crystal structures of the Uncleaved conformations for RBD Up (code: 6VYB) and RBD Down (code: 6VXX) were obtained from RCSB(6). These structures came with the 2P mutations, wherein the residues at K986 and V987 were replaced by Prolines (2P) to stabilize the trimer. The Furin cleavage was reinstated with residues 685-688 and reverted back from GSAS to the RRAR motif of the Spike protein. The glycosylation sites were modelled based on the most probable glycans on each site, obtained from mass spectroscopy data(13, 14).

Five independent production runs were then performed on AMBER using Charmm36m forcefield(15) for each system for a total run-time of 20 µs. For the analysis performed, the first 200 ns of all the trajectories were discarded to ensure that the systems had equilibrated (refer **SI Figure1**)

### Furin cleavage leads to stronger correlated motions across critical domains

To identify any differences in local fluctuations of the Cleaved and Uncleaved systems, we looked at the Root mean square fluctuation per residue (refer **SI Figure2**), which turned out to be similar except for the FCS region. For the Cleaved systems, the RMSF was much higher compared to the Uncleaved system due to the presence of an open loop. We decided to tease out cross-correlations between these observed local motions by calculating the Dynamic Cross Correlation Matrices (DCCM). DCCM is a method of representing time-correlated motions of each of the residues with all other residues throughout the protein[3]. Representing them in the form of a three-dimensional matrix, the plot shows that both the RBD Up and RBD Down conformations in the Cleaved system have significantly higher correlations than the Uncleaved system (see boxed regions of **Figure 1**). The higher correlations are especially observed in the regions of the NTD and RBD, which include both intra-chain and inter-chain interactions. We also see the reduced correlation between the Up RBD in Protomer B and NTD of Protomer C in the RBD Up conformation which indicates that the correlation is reduced upon the opening of the RBD. This is not surprising as the RBD moves further away from the neighbouring NTD when it undergoes the Down to Up transition. These observations were interesting, because contrary to our anticipation that due to the increased disorder at the FCS loop due to cleavage, there might be increased random motions in the Spike protein, thereby reducing correlation. However, we see that cleavage increases the correlated motions between different regions of the Spike protein and most importantly, the regions of NTD and RBD.

**Figure 1:**
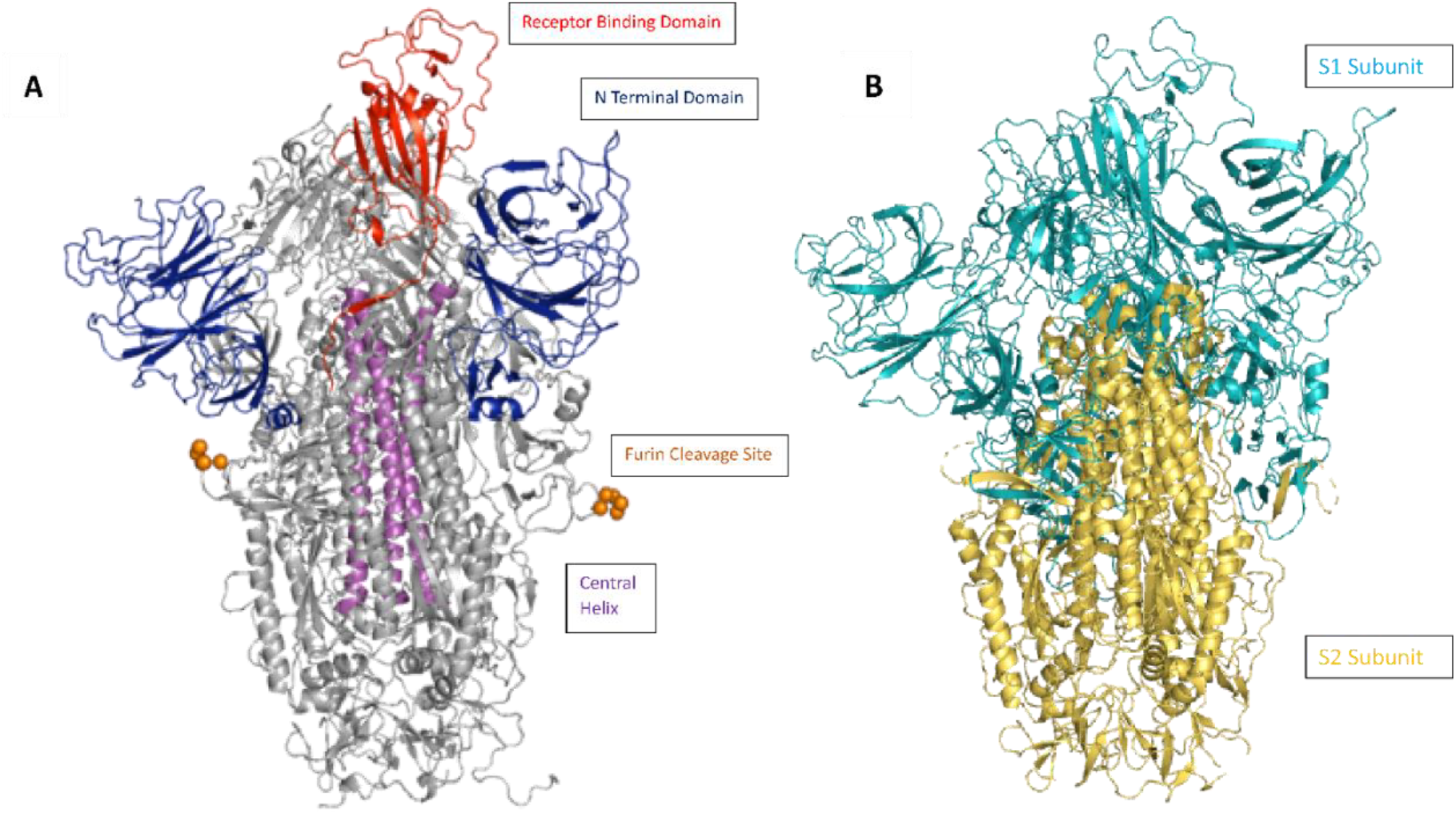
**Figure A** represents the key subunits of the Spike protein. The Up RBD is present on Protomer B. The Central Helix comprises of the bundle of helices present in the S2 domain. **Figure B** represents the two subunits of the Spike protein, S1 and S2.

**Figure 2.**
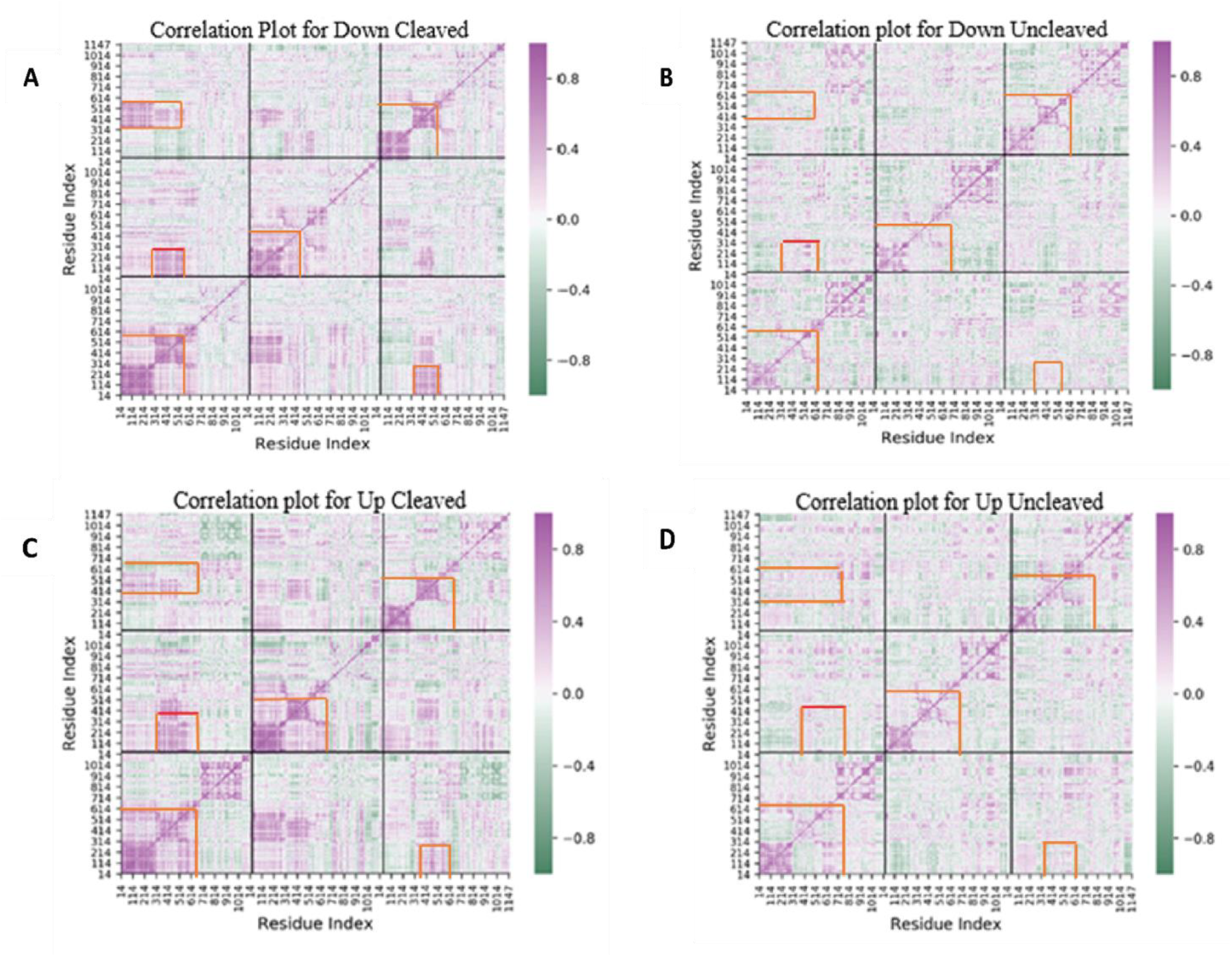
Cleaved conformation has a much higher correlation than the uncleaved especially observed in the NTD and RBD regions. **A, B** Cross-correlation plots for the Down Cleaved and Uncleaved conformations. **C, D** Cross correlation plots for the Up Cleaved and Uncleaved Conformations. The correlation is much higher in the Cleaved conformations in both Down and Up systems, especially in the regions of interest such as the NTD and the RBD (marked by red boxes), than the Uncleaved conformation

Based on our observations of these correlated motions, we wanted to check for differences in higher dimensional embedding. This was achieved using Principal Component Analysis (PCA), where we projected the eigenvectors of one conformation of the Cleaved system onto the corresponding conformation of the Uncleaved system. The key observation from PCA was that these systems sample completely different large amplitude pathways as indicated by the plots where the coordinate spaces are very different. It is interesting to note that the Cleaved conformation settles into basins much quicker than the Uncleaved conformation (check **SI Figure 2)** These differences in the essential motions between the Cleaved and Uncleaved systems, increased correlations in the NTD and RBD regions between the two systems make it clear that importance in antibody response and driving functionally relevant structural transitions makes these motions especially important.

### Furin Cleavage increases the probability of encountering ACE2 for binding

The Receptor Binding Domain (RBD) is responsible for binding to the ACE2 receptor and hence is the primary target for neutralizing antibodies (2). From our previous results, it was understood that the Cleaved conformation has a higher degree of correlated motions in the RBD and NTD for both within a protomer, as well as between different protomers. Hence, it is imperative that the RBD of these two conformations be studied in closer detail. The ability of the RBD to bind to ACE2 is highly dependent on its orientations and angle of tilt, as higher angles of outward tilt correspond to greater exposure, which translates to higher binding affinity (16). We quantified these motions by calculating the (i) tilt and (ii) torsional angles in two different cartesian directions. This data is represented in the form of Potential for Mean Force plots (PMF) 2D plots, where deeper basins correspond to higher stability.

When the RBD is Up, it can sample multiple tilt angles due to its higher degree of freedom. There are two orthogonal tilt angles possible: sideward and Up-Down. (see **SI Figure 4**). We observe that the FCS cleavage changes the conformational distribution of both these tilt angles. In the case of the up-down motion, when the FCS is intact, the RBD orientation is restricted in space. Upon cleavage of the FCS, the same RBD starts sampling varied degrees of up and down tilt angles. It is to be noted that the sideward tilt does not change the exposure of the Receptor binding motif, whereas the up-down motion can affect the ACE2 binding capability, varying between close-like and open-like conformations. Literature also suggests that RBD samples different conformations and our observation is when it is cleaved, it samples drastic changes in conformational space. Meanwhile, in the Uncleaved conformation, the RBD is more rigid in its tilt. These motions are represented in **Figures 4C** and **4D**. Hence it could be said that the Cleavage at Furin is enabling the RBD to sample a range of different tilts with the most dominant and stable ones being an inward tilt which is intrinsically stable and an outward tilt, which increases its probability of binding to an ACE-2 receptor(16)

**Figure 3.**
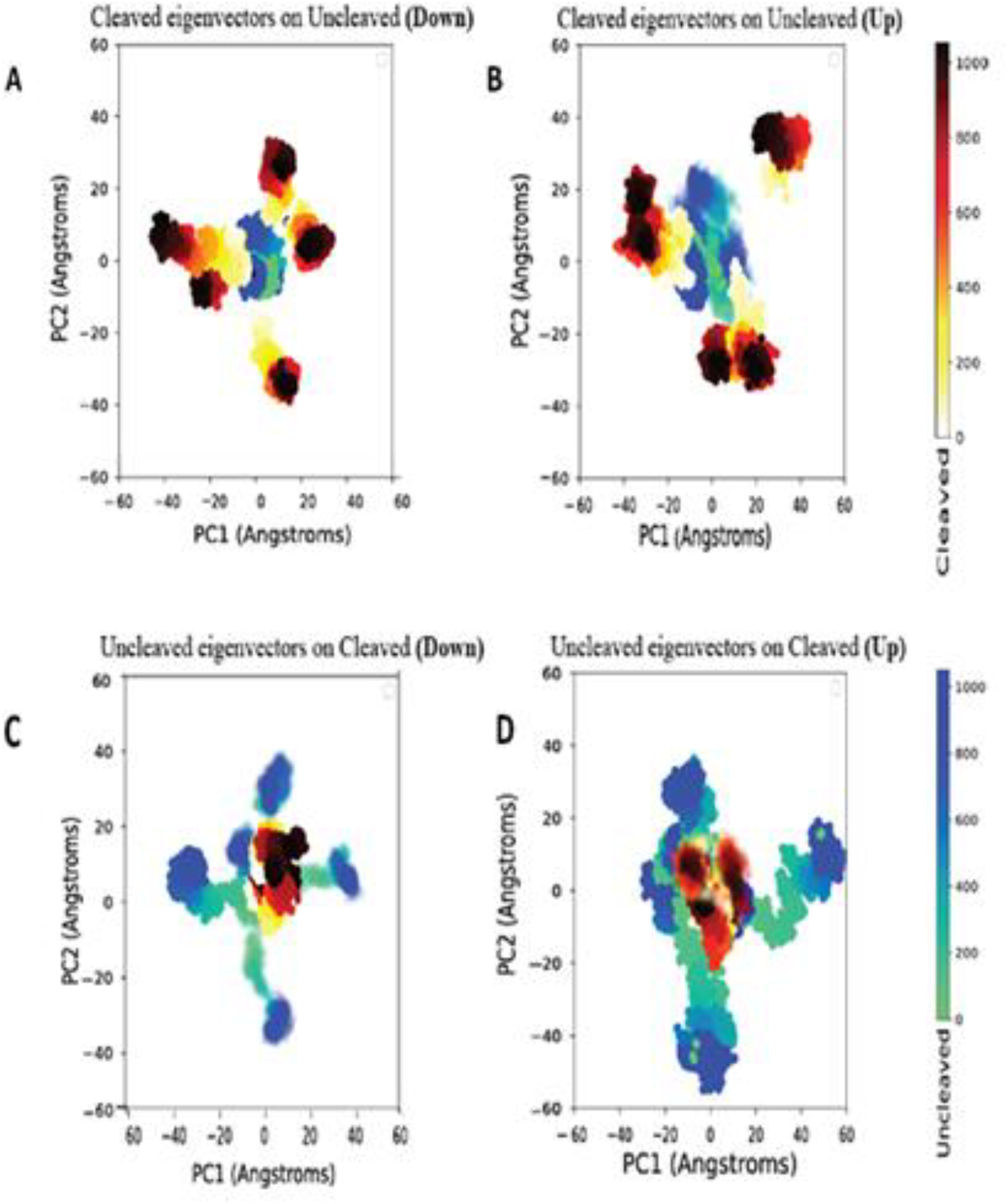
Furin cleavage of the Spike protein completely changes its large amplitude motions. **A**. PMF plot representing the Up-down bend of the RBD-Up conformation for both Cleaved and Uncleaved systems. **B**. PMF plot representing the sideward bend of the RBD-Up conformation for both Cleaved and Uncleaved systems. **C, D**. 3D representation of the corresponding bins in the Up-Down and sideward bend plots

**Figure 4.**
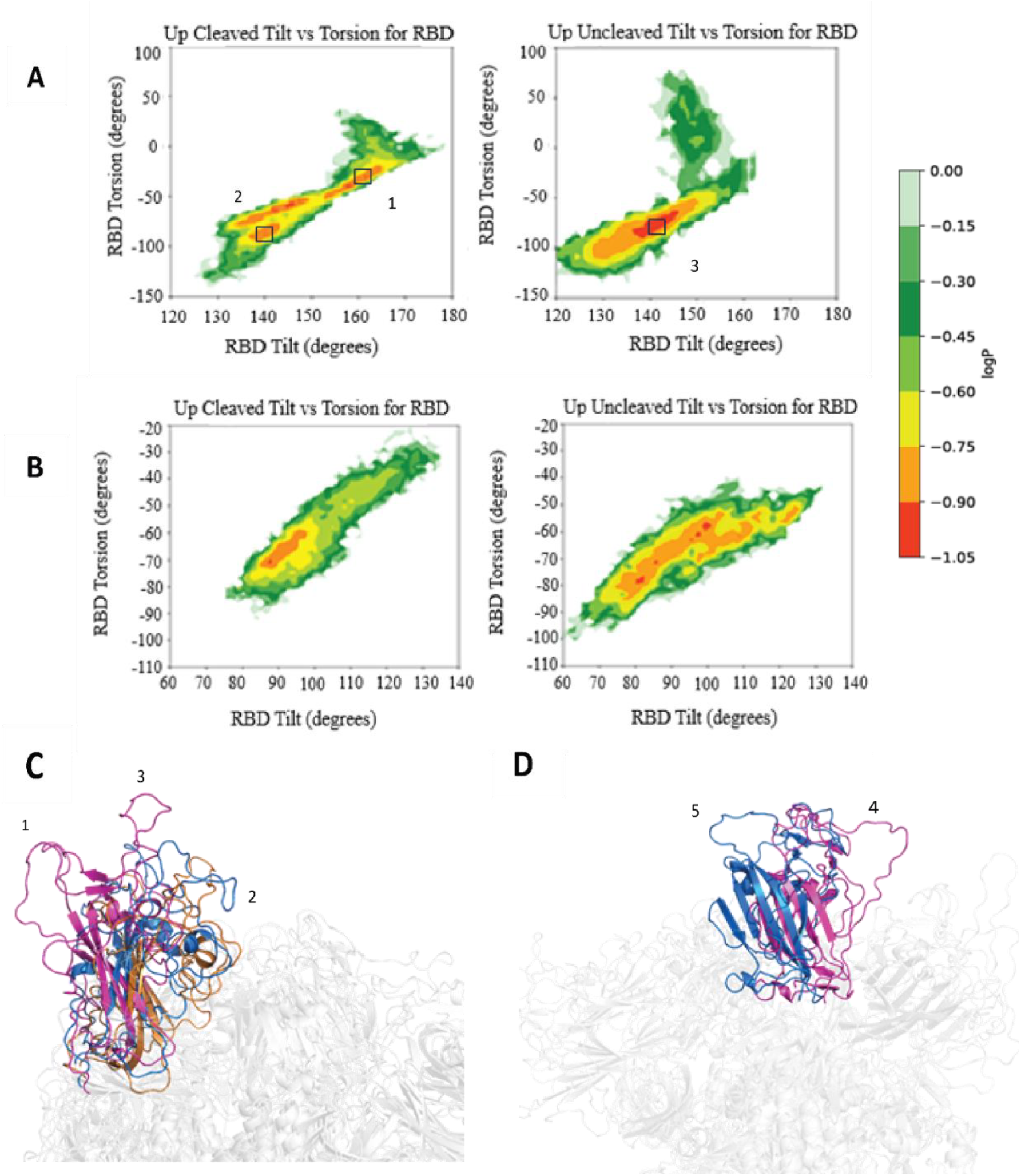
Furin cleavage of the Spike protein completely changes its large amplitude motions. **A**. PMF plot representing the Up-down bend of the RBD-Up conformation for both Cleaved and Uncleaved systems. **B**. PMF plot representing the sideward bend of the RBD-Up conformation for both Cleaved and Uncleaved systems. **C, D**. 3D representation of the corresponding bins in the Up-Down and sideward bend plots

### The motion of the NTD varies after the cleavage of Furin

It has been posited in the literature that the NTD’s motion greatly influences the RBD’s opening (17) and is also a key target for antibodies(18). NTD can either block the path of the RBD or move away, therefore acting as a wedge. Here too, we have represented these motions as PMF plots (**Figure 5**). NTD can either block the path of the RBD or move away, therefore acting as a wedge(17).

**Figure 5.**
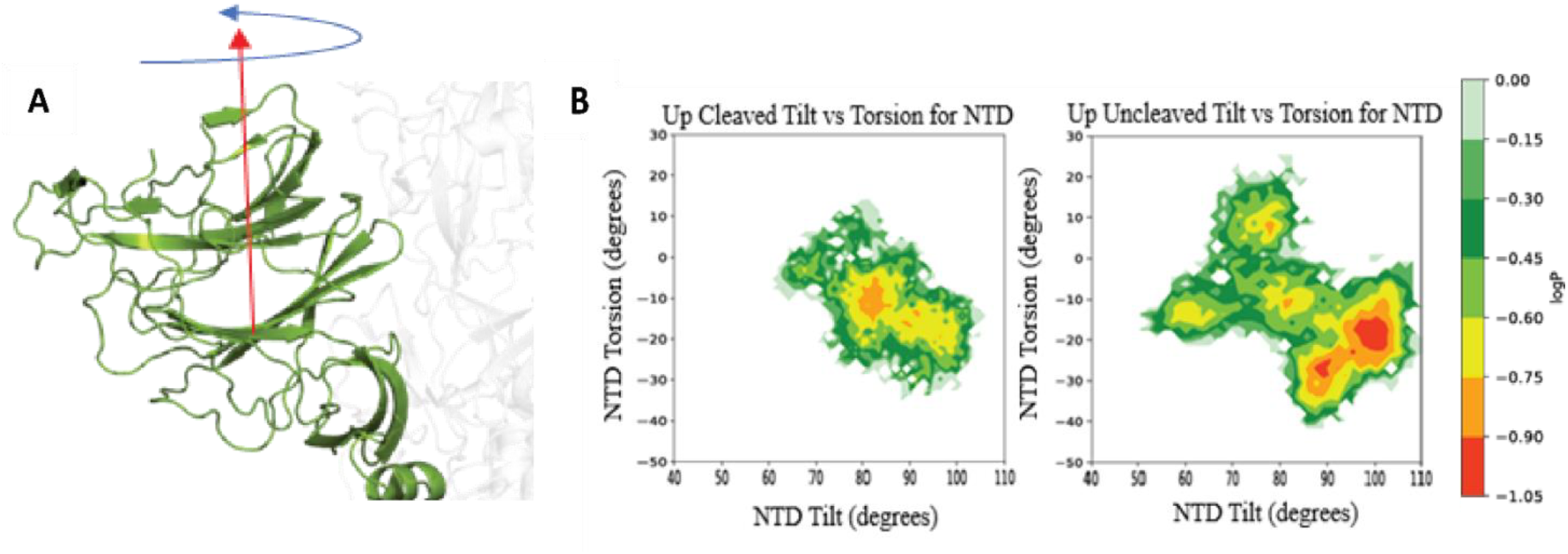
The NTD of the Up Uncleaved system samples a larger conformational space when compared to the Up Cleaved. **A, B**. These are the PMF plots for the Up Cleaved and Uncleaved conformations respectively. The NTD of the Uncleaved conformation samples a larger conformation space while the Cleaved conformation is more compact.

In our case for the RBD Down system, the tilt and torsional angles were plotted for all three NTDs combined as we wanted to look at the overall NTD motion. For both Down and Up systems in Cleaved, the NTD motions are more constrained. When the NTD is next to the Up RBD, the NTD is sampling a distributed landscape and its propensity to act as a wedge is much higher. There is less resistance to the opening of the RBD when there is cleavage

We observe that the NTD samples have different regions of stability for Cleaved and Uncleaved conformations. It is to be noted that the NTD samples regions of higher stability along all cartesian directions for the Cleaved conformation as compared to the Uncleaved conformation. This result agrees with our earlier observed DCC plots where the Down Cleaved conformation had a higher correlation in the NTD regions than the Down Uncleaved conformation.

As for the RBD Up system, the tilt and torsional angles were plotted for each NTD separately. We were especially interested in the motion of the NTD adjacent to the Up RBD as it is known to be responsible for the motion of the RBD(17, 18). We observed that the Uncleaved conformation samples a larger distribution of tilt and torsional angles, especially along the direction which correlates to the sideward swinging motion of the NTD. This motion is known to control the downward tilting motion of the RBD(17).

### The “Trunk” of the Cleaved conformation expands more than the Uncleaved

The S2 subunit of the Spike protein is responsible for viral fusion and entry into the host cell. It also contains two cleavage sites: 1) S1/S2 which is cleaved by Furin greatly enhances the capability for viral entry and is a precursor for viral fusion(19, 20) and 2) S2^I^ which is cleaved by TMPRSS and is responsible for membrane attachment and viral fusion(4, 21). As we saw earlier, the cleavage at S1/S2 brings about a huge difference in the overall dynamics of the system, most notably in regions such as RBD and NTD, making them more correlated. Cleavage at the Furin cleavage site also primes the Spike protein which enhances its ability for viral entry (22). This Furin-mediated pre-cleavage at the S1/S2 site in infected cells might promote subsequent TMPRSS2-dependent entry into target cells (20). We wanted to see if we could capture the effects of cleavage on the S2 domain which could lead to a cascade of events. Our understanding is that it is set in motion by S1 falling off, which releases spatial constraints. This exposes the FP region in the S2 subunit, which folds into a three-helix bundle along with the HR1 and CH(23). This elongated bundle formation allows the fusion peptide to insert into the host cell membrane. Our motive was to investigate if we could capture structural changes which could be precursors to these large-scale conformational transitions. This was done in two ways (**Figure 6**).

**Figure 6:**
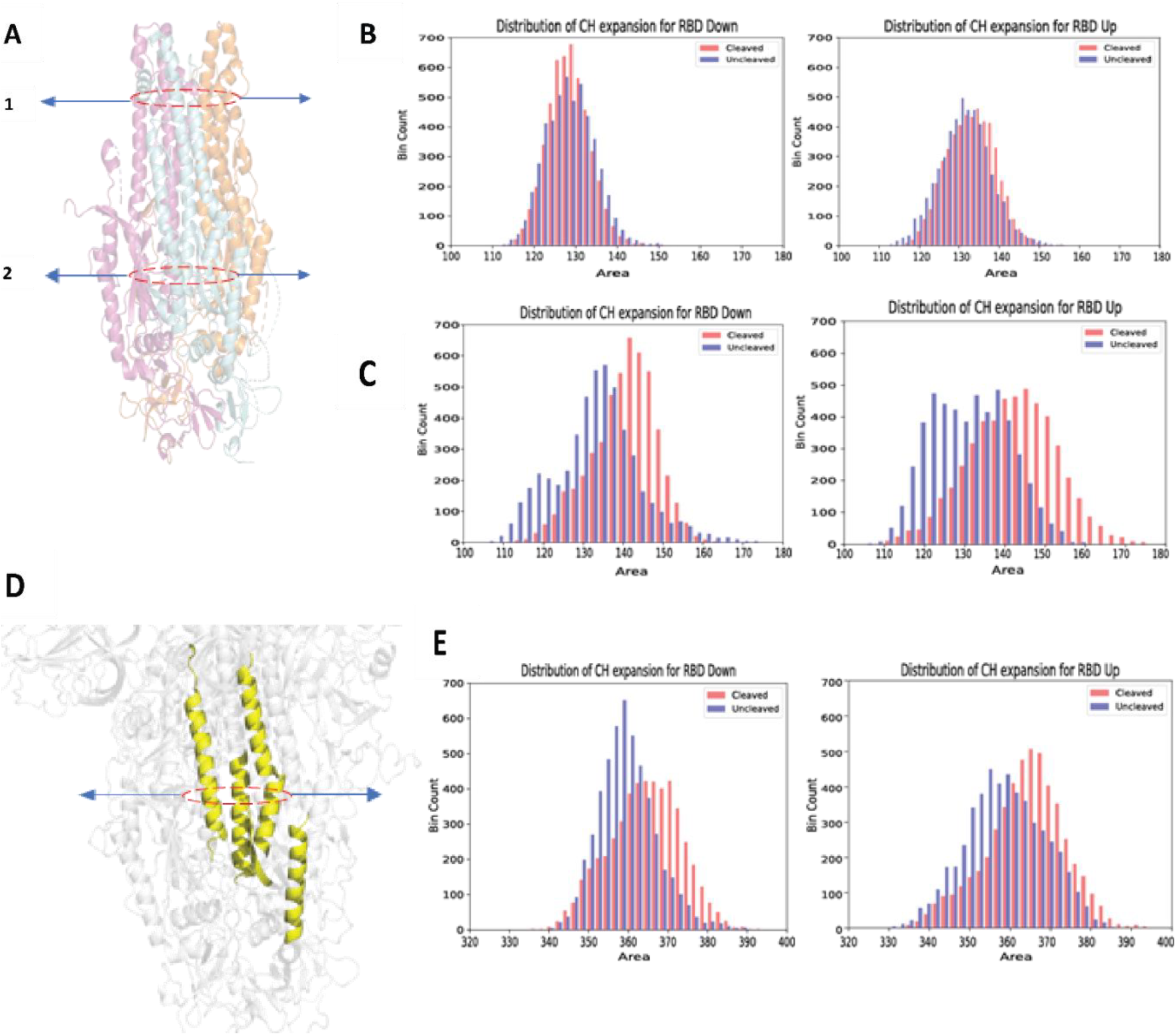
The S2 Subunit of the Furin cleaved Spike protein has a tendency to expand outwards. **Figures B** and **C** represent the distribution of areas of expansion of the Central Helix, as shown in **A1** and **A2**. It is seen that The Cleaved conformation for both the RBD Down and RBD Up shows an outward expansion of the Central Helix. These results are further confirmed by calculating the expansion of the Center of Mass of the Central Helix as shown in **Figure D**. We do observe that the Cleaved conformation has a higher degree of expansion when compared to the Uncleaved conformation as shown in **Figure E**.

Firstly, we looked at the “opening” of the S2 Subunit at two different locations of the Central Helix, namely at residues 1000 and 1030 for both the RBD-Up and Down conformations of the Cleaved and Uncleaved systems. This was measured by looking at the area of expansion of these residues and comparing the Cleaved and Uncleaved conformations. We find that the Cleaved system for both RBD Up and Down conformations has a higher area of expansion than the Uncleaved system. This indicates that the lower portion of the Central Helix undergoes a certain outward expansion upon Furin Cleavage. These results were further confirmed by looking at the expansion of the Center of Mass of the Central Helix. This was calculated for helices in each protomer, and the area of expansion was measured. This also proved that the Cleaved systems have a higher area of expansion in the Central Helix region.

To obtain a quantitative understanding of how different the distributions of areas are, we performed a Kolmogorov-Smirnov test (K-S)(24). The K-S test statistics give an estimate as to how similar or dissimilar the two distributions are. The test results proved that the two distributions we see are indeed statistically significantly different, with the Cleaved system having a distribution of higher areas of expansion. This could lend weight to the fact that Furin Cleavage of the Spike protein primes it and makes it easier for the Spike protein to undergo further downstream processing steps such as cleavage at S2^1^.

### Glycans are clustered in the Cleaved conformation especially around the RBD and NTD

So far in our analysis, we have looked at the protein dynamics of our system and we observed quite a lot of differences between Cleaved and Uncleaved systems. Since our systems are fully glycosylated, we must look at glycans as it is known that they influence protein stability(25) and play a crucial role in viral dynamics and pathogenesis(26). The Spike protein is heavily glycosylated and has around 22 N-glycosylation sites and 17 O-glycosylation sites(27). Glycans on the NTD, especially on residues N165 and N234, are known to be responsible for controlling the motion of the RBD(26, 28).

Our idea was to see if the glycans contributed to the differences observed between the Cleaved and Uncleaved conformations. This was done by extracting the structures which were in the stable basins as observed in the PMF plots (**Figure 7**). Our areas of interest were the NTD and RBD and as we previously mentioned the glycans around those subunits control the motion of the RBD. We observed that the glycans seemed to have a higher degree of clustering in the Cleaved conformation when compared to the Uncleaved conformation. This was especially observed in glycans N165 and N234 which, as mentioned earlier, control the RBD motion. It could be hypothesized that this could be a factor in the observed differences in the RBD motions between the Cleaved and Uncleaved conformations.

**Figure 7.**
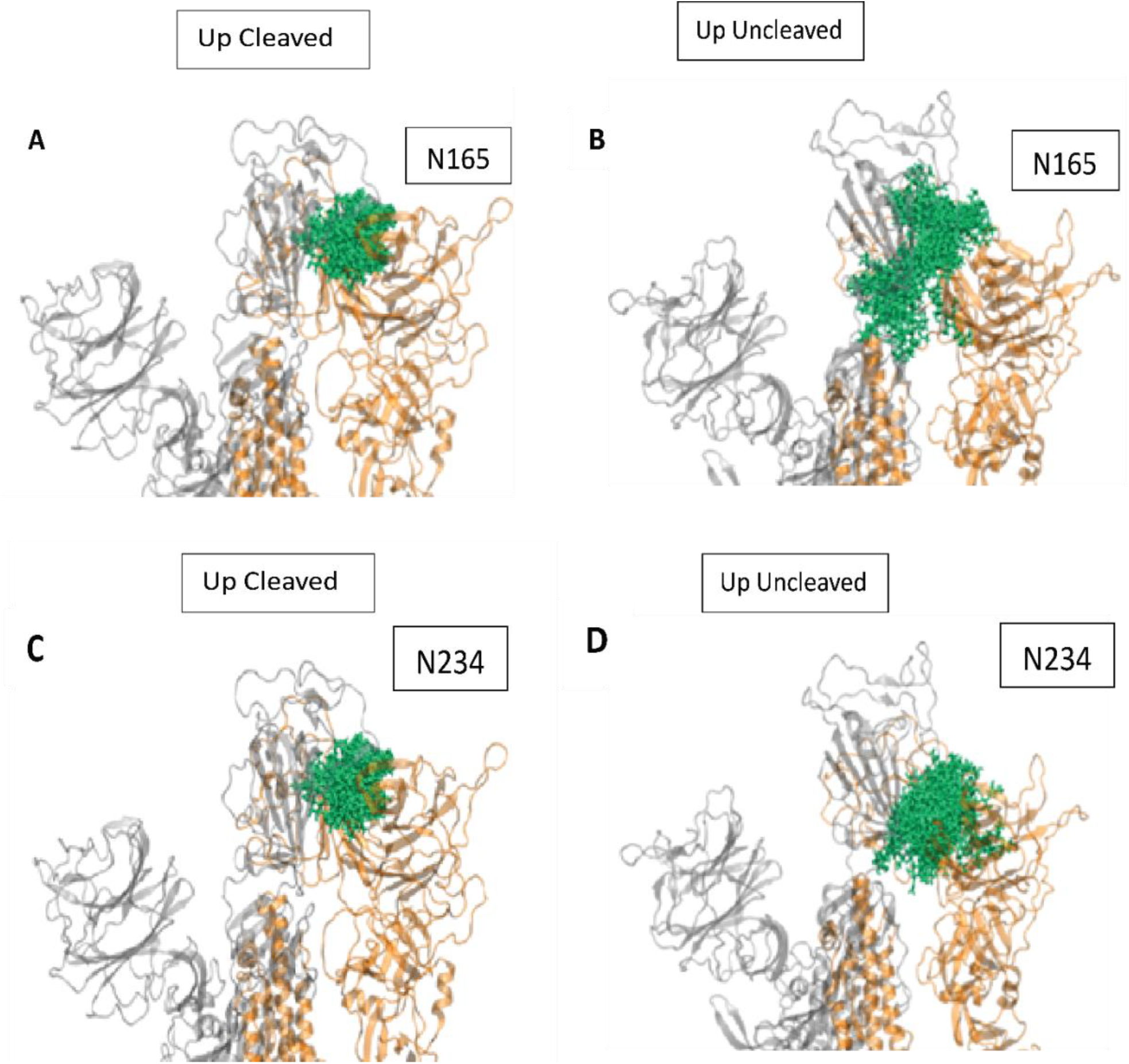
Glycans appear to be highly clustered in the Cleaved conformation when compared to the Uncleaved conformation which could be responsible for the observed dynamics of the NTD and the RBD. **A, B** The glycans on N165, **C, D** The glycans on N234

## Discussion

There has been debate regarding whether Furin cleavage occurs before or after ACE-2 binding (29). The commonly used structure for research and current design of vaccines based on the Spike protein have the Furin cleavage site (FCS) mutated to GSAS which keeps the Spike protein intact(7). However, recent literature seems to suggest that the cleavage occurs during the virus particle formation itself, (8, 19) as there is an abundance of Furin and other proteases inside lung cells. It has also been suggested that the cleavage at the S1/S2 subunit confers useful properties to the Spike protein. It destabilizes the Spike protein which enhances its ability to bind to ACE-2(8-10). It also helps the down conformation to open thus making it ACE-2 accessible(7, 10). It also makes it easier for the S2 cleavage by TMPRSS(22, 30). As previously discussed, the deletion of the RRAR motif or its mutation to GSAS in experimental studies does not accurately represent the physiological nature of the Spike protein as the motif is cleaved during the viral formation itself. As a result of these properties, it is important to study the post-cleavage dynamics of the Spike protein to design effective therapeutic strategies

The Omicron strain has been detected with the highest number of mutation sites(31). There have been around 15 mutations in the RBD, closer to the ACE-2 binding interface, for the Omicron strain when compared to only two in the Delta strain(31). It is possible that the presence of these additional mutations caused the increased transmissibility of the virus. While the WHO has no currently designated Variants of Concern (VOCs) circulating, BA.2.86 and JN.1, which are the subspecies of the Omicron strain, are the current Variants of Interest (VOI). The BA.2.86 strain has 30 mutations in the Spike protein, which is equivalent to the number of mutations in the original strain of Omicron from the Wuhan strain. Since we wanted to look at the impact of cleavage on protein allostery, our interest was in teasing out the differences in the large amplitude motions between the pre-cleavage and post-cleavage conformations of the Spike protein. Our initial assessment was to look at the local fluctuations by calculating the Root mean square fluctuation (RMSF), which was very similar between the Cleaved and Uncleaved systems except at the FCS, where it was higher for the Cleaved conformation. We decided to tease out cross-correlations between these observed local motions by calculating the Dynamic Cross Correlation Matrices (DCCM). The analysis showed that both the RBD-Up and RBD-Down conformations in the Cleaved system have significantly higher correlations than the Uncleaved system. Higher correlations are especially observed in the regions of the NTD and RBD, which include both intra-chain and inter-chain interactions. Based on our observations of these correlated motions, we proposed to check for differences in higher dimensional embedding using techniques such as Principal Component Analysis (PCA). This led to the observation that the Cleaved and Uncleaved systems sample completely different dynamics.

Since it has now been established that there are clear differences in the motions of the NTD and RBD between the Cleaved and Uncleaved conformations, as evinced by our PCA and DCC plots, we looked at the individual motions of the RBD and NTD. This was achieved by estimating the tilt and torsion of the NTD and RBD in three different directions and calculating the PMF values to find regions of structural stability. These motions are especially important as the binding affinity of ACE-2 depends on the angle of the RBD, larger angles lead to higher binding affinity. (16). The motion of the NTD, especially that which is adjacent to the RBD, plays a significant role in the motion of the RBD. It can either tilt outwards which helps the RBD to tilt downwards which is an intrinsically stable conformation or it can act as a wedge which then prevents the RBD from falling completely(17).

For the Up Cleaved System, we find that the RBD samples two basins of higher stability which correspond to low and high tilt angles. This indicates that the RBD moves outwards and inwards, both of which are stable. The outward motion enables the RBD to bind ACE-2 more effectively whereas the inward motion, although reduces the binding affinity, might help it evade antibodies. This motion is done in tandem with the NTD as we observe the wedging motion which prevents the RBD from falling. The higher degree of stable conformational sampling of the RBD in the Up Cleaved conformation could indicate an effort to increase its probability of binding to ACE-2, whereas the Uncleaved conformation is more rigid. These findings find agreement in experimental literature where it was observed that the Cleaved Spike protein is destabilized and has a greater propensity to adopt an “open” conformation than its Uncleaved counterpart(10).

As mentioned earlier, the cleavage at the Furin cleavage site primes the Spike protein which enhances its ability for viral entry and also promotes S2 cleavage by TMPRSS2(22). We wanted to check if there was any effect of Furin cleavage on the S2 subunit. This was initially checked by calculating the area of expansion of the Central Helix (CH) which revealed that the Cleaved conformation has a higher degree of expansion in the S2 region than the Uncleaved. To further consolidate our findings, we calculated the motion of the Center of Mass (COM) of the helices present in the S2 domain, which also proved that the Cleaved conformation has a higher degree of expansion in the S2 domain. This could mean that the Furin cleavage primed the Spike protein, which makes it easier to undergo further processing. The correlated motions observed in the RBD and NTD regions might have an allosteric effect on the observed priming.

It is important to discuss the role of glycans in this study as they contribute to viral effectiveness(26) and also influence protein dynamics and stability(25). In the context of the SARS Cov-2 Spike protein, it has been established that glycans N165 and N234 heavily modulate the motion of the RBD, especially the ‘open’ conformation and provide extensive(26, 28). Glycan N343 on the RBD plays a key role in the interaction with ACE-2 receptor and acts as a “gate” in pushing the RBD from the “closed” to an “open” conformation(28, 32). These findings prompted us to check similar motions in our Spike protein and to see how they differ between the Cleaved and Uncleaved systems. We find that the glycans of interest in the NTD and RBD take up stable conformations like the protein. This could indicate that the conformational stability of the protein influences the glycan’s conformational stability and vice-versa. Glycans N165 and N234 on the NTD in the Cleaved conformation also seem to act like a wedge which controls the tilting of the RBD.

Our findings indicate that the RRAR cleavage site plays an important role in the pathogenesis of the virus. The cleavage by Furin induces several large amplitude motions in the Spike protein, especially in the RBD, which improved the chances of ACE2 binding. The cleavage also helps in “priming” the S2 domain of the Spike protein which could help further cleavage at the S2^1^ domain. The role of glycans was also studied with the glycans of interest being N165 and N234 on the NTD which helps in controlling the motion of the RBD, and N343 on the RBD which contributed to shielding. Hence these results provide a quantitative framework that allows us to capture the impact of Furin cleavage on the Spike protein and help us to design effective therapeutic strategies. This study can also be extended to other Betacoronaviruses such as MERS and viruses which possess a Furin Cleavage Site such as HIV and Influenza.

## Funding and Acknowledgements

N.M., R.S. and S.C. were supported by NIH NIGMS grant grant R35GM151231-01, and through NEU Faculty Startup Funds. AI158571. This research used computational resources from Northeastern Discovery cluster at MGHPCC, as well as through NSF ACCESS.

## Author contributions

N.M. and S.C. conceptualized the study and designed the experiments. S.C. prepared the structures. N.M. ran the simulations. N.M. and R.S. performed the data analysis. N.M. and S.C. performed data interpretation. N.M. prepared the figures. N.M and S.C wrote and edited the manuscript. S.C. is the corresponding author.

## Competing interests

The authors declare no competing interests.

## Data and materials availability

Data and materials are available from the corresponding authors upon request.

## Notes

### Competing Interest Statement

The authors have declared no competing interest.

