## Supplementary material for "Cleaved vs. Uncleaved: How Furin Cleavage Reshapes the Conformational Landscape of SARS-CoV-2 Spike": Methods and Supplemental Information

**Model setup:** The crystal structures of the Uncleaved conformations for RBD-Up (code: 6VYB) and RBD-Down (code: 6VXX) were obtained from RCSB(1) which were the starting structures. The Furin cleavage was reinstated by reverting the modified GSAS to the RRAR motif. We also induce cleavage at the FCS by introducing a "cut" in the initial structure. Additionally, all the starting structures included a 2P mutation wherein, the residues at K986 and V987 were replaced by Prolines (2P) to stabilize the trimer. To accurately mimic the physiological behavior of the Spike protein, the most probable glycan types at each glycosylation site were selected based on mass spectrometry results and added by ab initio modelling using ALLOSMOD(2) with CHARMM36 forcefield. All visualizations and structural representations were performed with VMD 1.9.3(3).

**Simulation Protocol:** We have a total of 4 distinct systems: Furin Cleaved Spike protein with the RBD Up (referred to as **Up Cleaved**), Furin Cleaved Spike protein with the RBD Down (referred to as **Down Cleaved)**, Furin Uncleaved Spike protein with the RBD Up (referred to as **Up Uncleaved)** and Furin Uncleaved Spike protein with the RBD Down (referred to as **Down Uncleaved).** A TIP3P water model was used for simulations and a cubic waterbox was used as the simulation box with a padding of 20 $A^0$ to account for the motion of glycans. The systems were neutralized at an ionic concentration of 150 mM of KCl. For all these systems, steepest decent energy minimization was performed with a gradual release of constraints on the glycan-heavy atoms and protein backbone. The first equilibration step involved a restrained simulation in a constant volume and temperature (NVT) ensemble for 2 ns. The second step involved a restrained simulation with constant pressure and temperature (NPT) for 10 ns. During the first and second stages, harmonic position restraints were imposed on all non-hydrogen atoms. The third step of equilibration involved 10 ns of simulation in the NPT ensemble and only included position restraints on protein backbone atoms.

We ran 5 separate trajectories for each system, each around 1.2μs in length. We discarded the initial 200ns of each trajectory to account for stabilization, using the Berendsen barostat, the particle mesh Ewald method for electrostatics and temperature coupling at 310K with a Langevin thermostat. Hence the total runtime was 5μs for each system. In total, 20 simulations were performed for an aggregated time of 20 $\mu$s. Since we had access to multiple GPUs on the High-Performance Computing (HPC) cluster, we converted our input files to a format compatible with the AMBER simulation package(4), which provides the best throughput on GPUs for our simulations. We also decided to use the Hydrogen mass repartitioning feature in AMBER which helps to speed up the simulations by using a timestep of 4fs.

**Root mean square fluctuations (RMSF):** RMSF is a measure of how much individual atoms in a molecule fluctuate from their average position over time. It is calculated for an atom *i* as:

$$RMSF_i = \sqrt{\frac{1}{T}\sum_{t_j=1}^{T}\left\|\boldsymbol{r}_{\mathrm{i}}(\mathrm{t_j}) - \boldsymbol{r}_i{}^{\mathbf{ref}}\right\|^2}$$

where $T$ is the total time of the production run, $t_j$ is the index for the time point of calculation, $\boldsymbol{r}_i^{ref}$ is the coordinates of the mean structure, and $\boldsymbol{r}_i$ is the coordinates of the $i^{th}$ frame. These calculations were performed using the MDTraj calculations(5) and are shown in **SI Figure 2.**

**Principal component analysis (PCA):** PCA is a dimensionality reduction technique, which in the case of MD simulations, can be used to extract dominant modes of structural variations and identify clusters with the trajectory. PCA was performed on all 4 systems: Up Cleaved, Down Cleaved, Up Uncleaved and Down Uncleaved for the Cα atoms. The Covariance matrix was calculated as:

$$C_{nn'} = \frac{1}{M} \sum_{m=1}^{M} (\vec{r}_{mn} - \langle \vec{r}_n \rangle) \otimes (\vec{r}_{mn'} - \langle \vec{r}_{n'} \rangle)$$

Diagonalizing this covariance matrix gives the eigenvectors describing the directions of the highest change in the amplitude of protein motion that translated to a conformational shift within the trajectory. We analyzed the top two components along which we see the highest variance. While individual PCA plots were interesting, we found that by projecting the eigenvectors of one system on the other, we were able to elicit differences in motions (see **Figure 3** and **SI Figure 3**). These calculations were performed using the 'covar' and 'anaeig' commands of GROMACS(6) along with the PCA module of Scikit-learn(7) and MDtraj (5).

**Dynamic Cross-Correlation (DCC):** DCC can provide insights into the correlative motions of atoms in a trajectory. This correlative motion between two atoms *i* and *j* is defined as

$$DCC(i,j) = \frac{\langle \Delta \mathbf{r}_i(t) \cdot \Delta \mathbf{r}_j(t) \rangle_t}{\sqrt{\langle \|\Delta \mathbf{r}_i(t)\|^2 \rangle_t} \sqrt{\langle \|\Delta \mathbf{r}_j(t)\|^2 \rangle_t}},$$

where $r_i(t)$ denotes the vector of the $i^{th}$ atom's coordinates as a function of time t, $\langle\ \rangle_t$ is the time ensemble average and $\Delta r_i(t) = r_i(t) - \langle r_i(t)T \rangle_t$ (8). The values are then plotted in the form of a NxN heatmap, where N is the number of Cα atoms in the system. The correlation values are calculated between -1 and 1, where 1=complete correlation; -1=complete anti-correlation; 0= no correlation. The DCC matrix was generated using the MD-Task Python library(9) and plotted using MatPlotLib(10). This was done for all 4 systems and is shown in **Figure 2**.

**Coordinates for studying RBD motion:** To describe the motion of the RBD, we define a downward bending and twisting motion of the RBD (ref **SI Figure 4**), which are measured relative to the central helix (CH). A vector representing the RBD spanning residuesY508-Cα and S514-Cα and the CH (reference vector) spanning residues I770-Cα and Q1010-Cα are defined. The bending of the RBD is then calculated as the angle or dot product between the reference vector and the RBD vector. The twist of RBD is defined as the dihedral angle that is formed by I770-Cα and Q1010-Cα of the reference vector, and Y508-Cα and S514-Cα of the RBD vector.

Another direction of RBD motion, which describes the sideward bend and twisting motion (ref **SI Figure 4B**), is also defined relative to the CH. The vector spanning the RBD is along D398-Cα and I434-Cα while that of the CH is I770-Cα and Q1010-Cα. The sideward bend angle is the dot product of these two vectors while the twisting motion is the dihedral angle between D398-Cα and I434-Cα, and I770-Cα and Q1010-Cα.

**Coordinates for studying NTD motion:** To describe the motion of the NTD, we define a sideward swinging and twisting motion (ref **SI Figure 5**), which are measured relative to the central helix (CH). A vector representing the RBD spanning residues I128-Cα and P225-Cα and the CH (reference vector) spanning residues Q1005-Cα and A1016-Cα are defined. The sideward swinging of the NTD is then calculated as the angle or dot product between the reference vector and the NTD vector. The twist of NTD is defined as the dihedral angle that is formed by Q1005-Cα and A1016-Cα of the reference vector, and I128-Cα and P225-Cα of the NTD vector.

This particular analysis is done for the NTD adjacent to the RBD for the Up systems (Cleaved and Uncleaved) as it has an impact on RBD motion(11). The other cartesian directions are also calculated (see **SI Figure 5**). A similar analysis is also done for the Down systems (Cleaved and Uncleaved); however, the average value is taken for all three NTD's for each cartesian direction (see **SI Figure 3**)

**Describing the expansion of the CH domain:** To describe the expansion of the CH domain, we employ two techniques. We first look at the area of expansion at two different regions of the stalk, namely at residues 1000 (further up the stalk) and residue 1030 (further below, closer to the S2[1] cleavage site). The area of expansion was approximated as the area formed by the triangle at those two residues across the three chains. Secondly, to confirm the results from above, we also looked at the expansion of the Center of Mass (COM) of the helices present in the CH domain. The COM of the bundle of helices was calculated for each chain and the area of expansion was again approximated as the area formed by the triangle.

To understand if the differences in these expansions for the Cleaved and Uncleaved conformations were statistically significantly different, we used the two-sample Kolmogorov Smirnov (KS) test. This test compares two independent samples to determine if they come from the same underlying distribution. It is based on the maximum absolute difference between the empirical cumulative distribution functions (ECDFs) of the two samples given by the formula:

$$D_n = \sup_x |F_1(x) - F_2(x)|$$

where $F_1(x)$ and $F_2(x)$ are the ECDFs of the two samples. The null hypothesis ($H_0$) states that the two samples follow the same distribution, while the alternative hypothesis ($H_1$) suggests they differ. A large $D_n$ value, with significance determined using KS distribution tables, indicates a deviation from $H_0$ meaning that the two distributions are statistically significantly different.

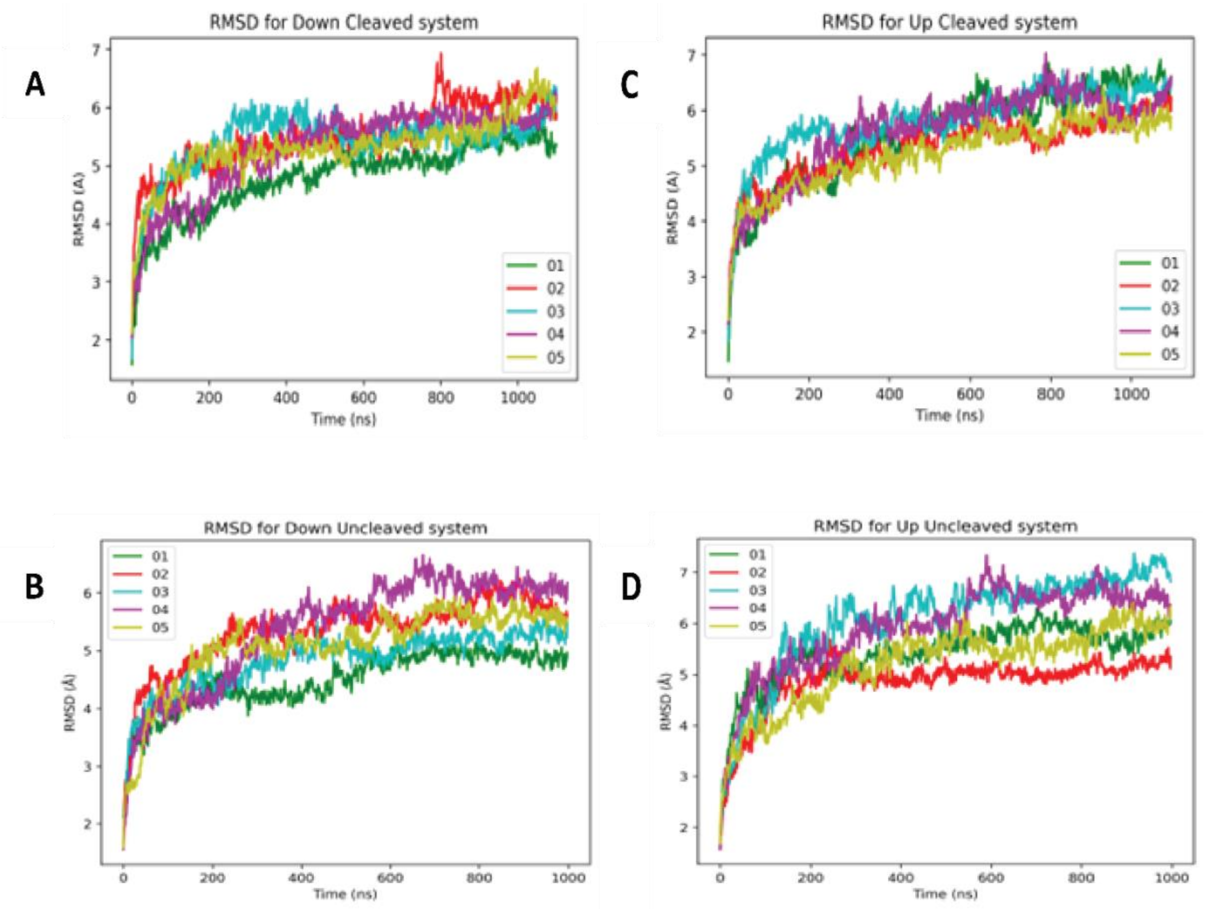


**SI Figure 1.** RMSD plots for the **A, B.** RBD Down and RBD Up of the Cleaved conformations and **C, D.** RBD Down and RBD Up of the Uncleaved conformations.

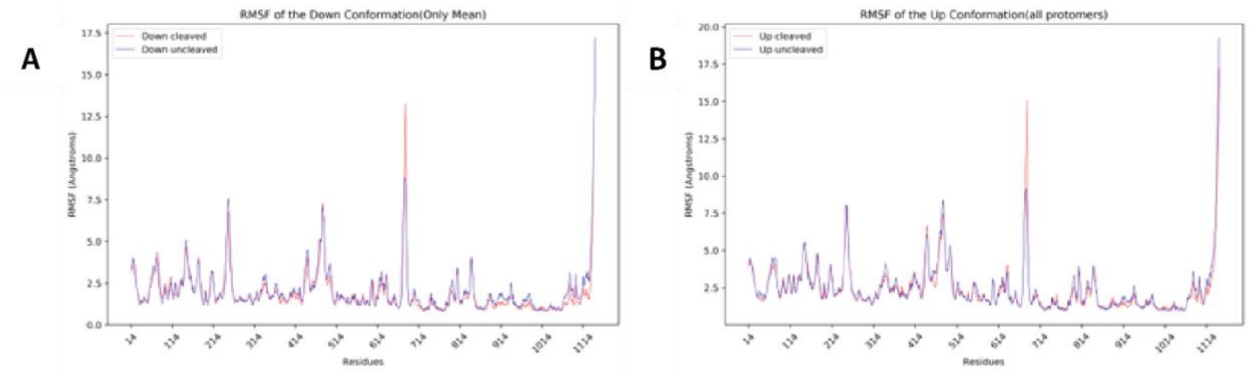


**SI Figure 2.** There is no difference between the local motions of the Cleaved and Uncleaved conformations. **Figures A** and **B** both show no difference in the local fluctuations between the Cleaved and Uncleaved conformations for both RBD Up and RBD Down except at the FCS region.

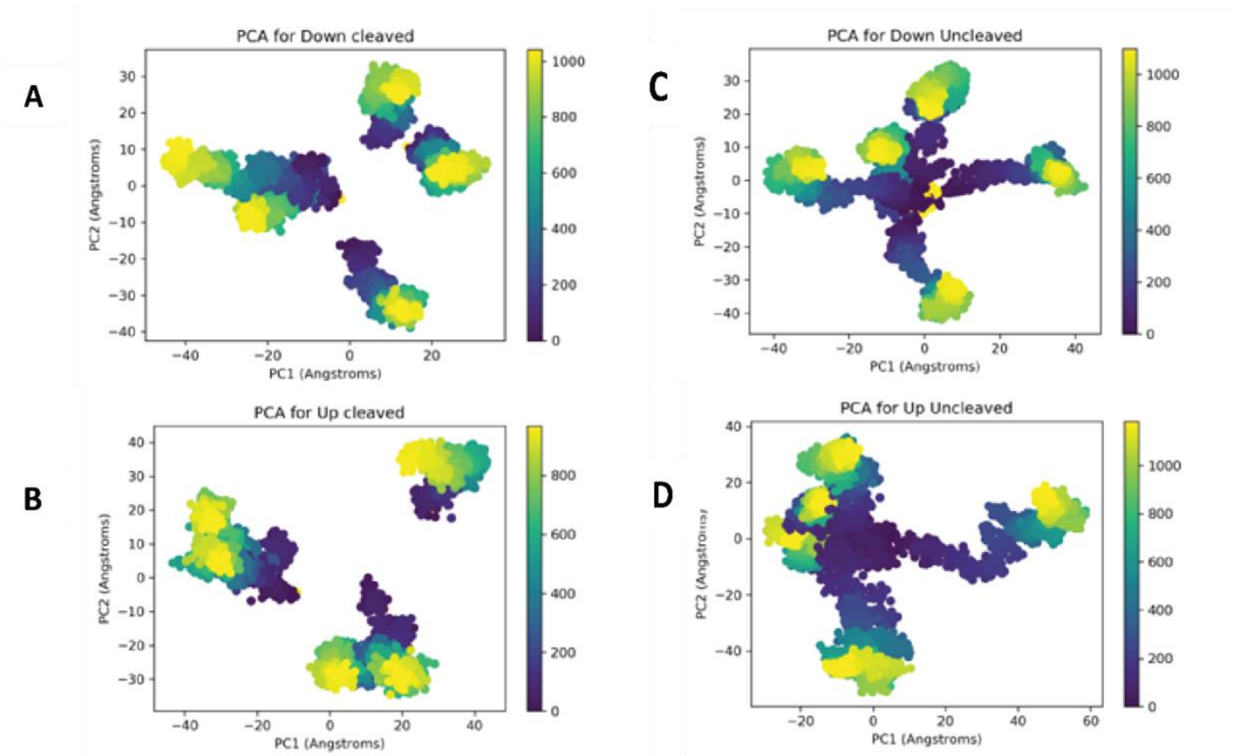


**SI Figure 3.** The Cleaved conformations for both RBD Down and RBD Up settle into basins faster than the Uncleaved conformations. **Figures A** and **C** represent the PCA for the Cleaved conformation and it can be observed that they reach their stable conformations much faster than the Uncleaved conformations in **B** and **D**

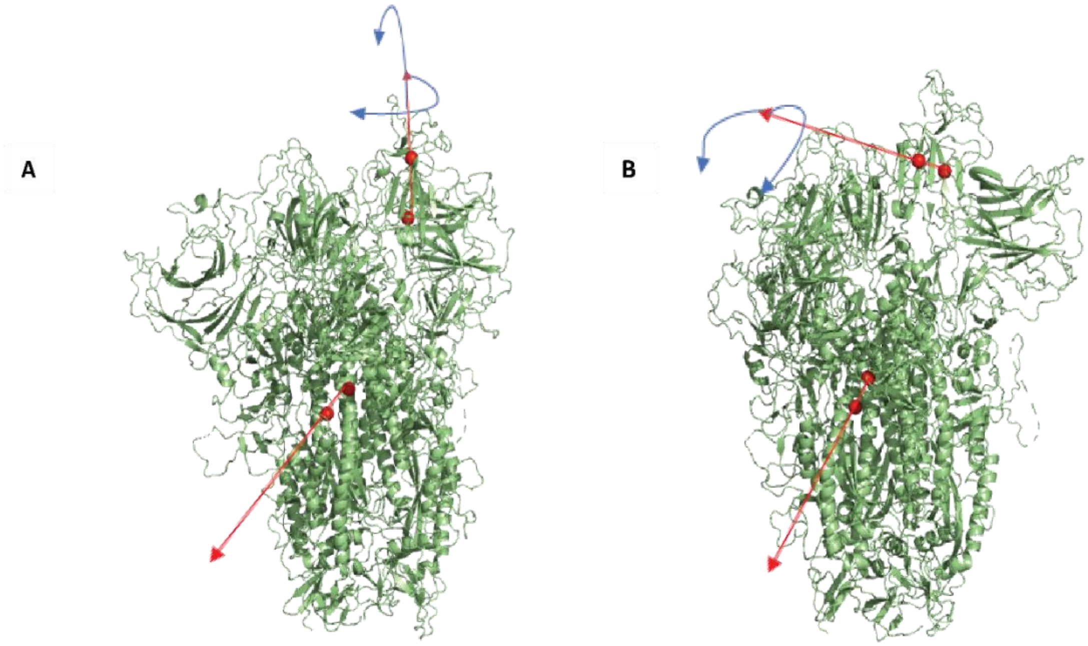


**SI Figure 4.** The coordinates for RBD motion are shown here. Figure **A** represents the downward bending motion of the RBD while **Figure B** represents the sideward tilt motion.

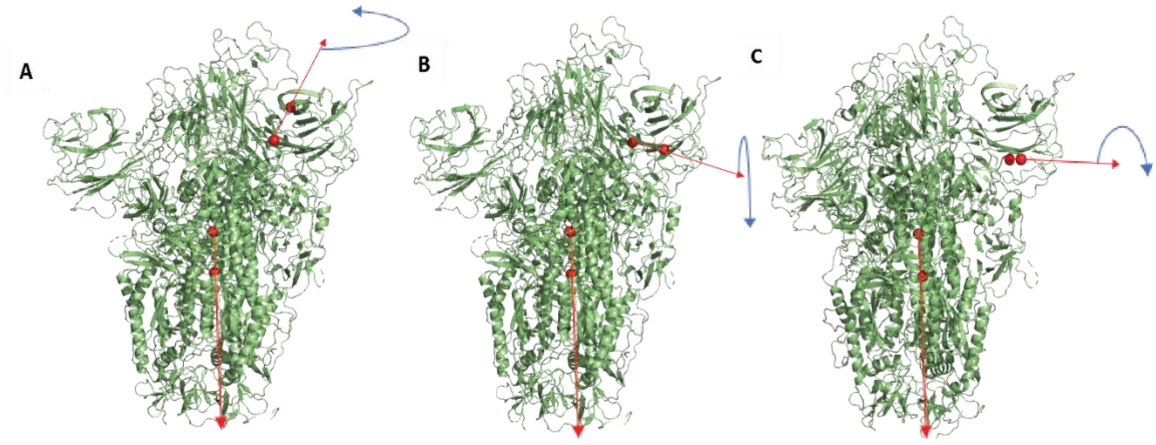


**SI Figure 5.** The coordinates for NTD motion are shown here. Similar residues are selected for the Down systems. **Figure B** is the swinging motion of the NTD which was discussed in the earlier sections.

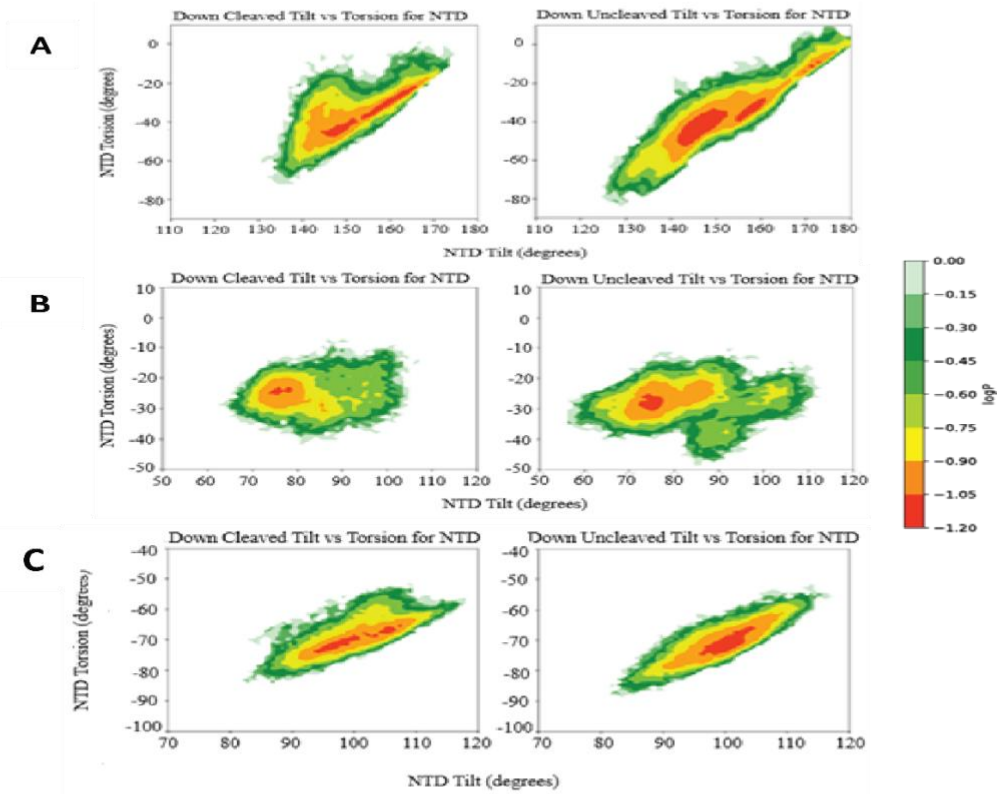


**SI Figure 6.** PMF plots representing the distribution of energy basins along different cartesian directions where the energy values represent the average for all NTD's. The directions are in the same order as in **SI Figure 5.**

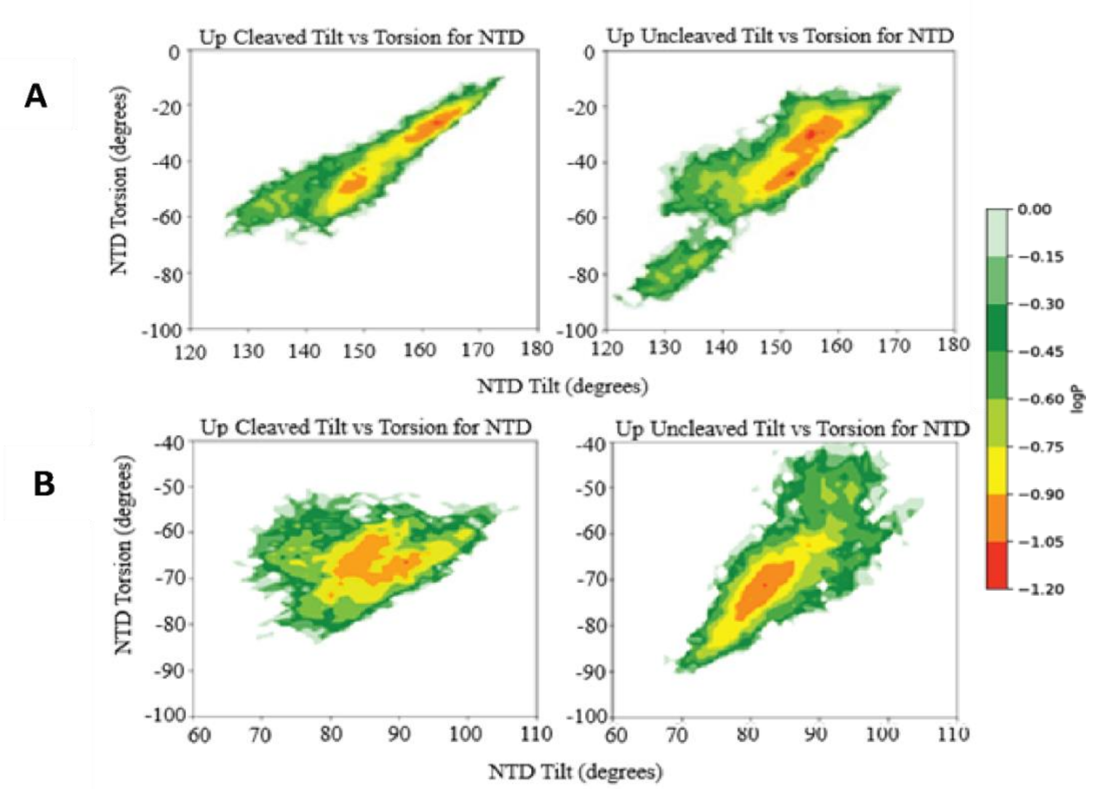


**SI Figure 7.** PMF plots representing the distribution of energy basins along the other two different cartesian directions for the NTD on protomer B. The cartesian directions are shown in **SI Figure 1** and **SI Figure 3**. The direction along **SI Figure 2** is discussed in the previous sections

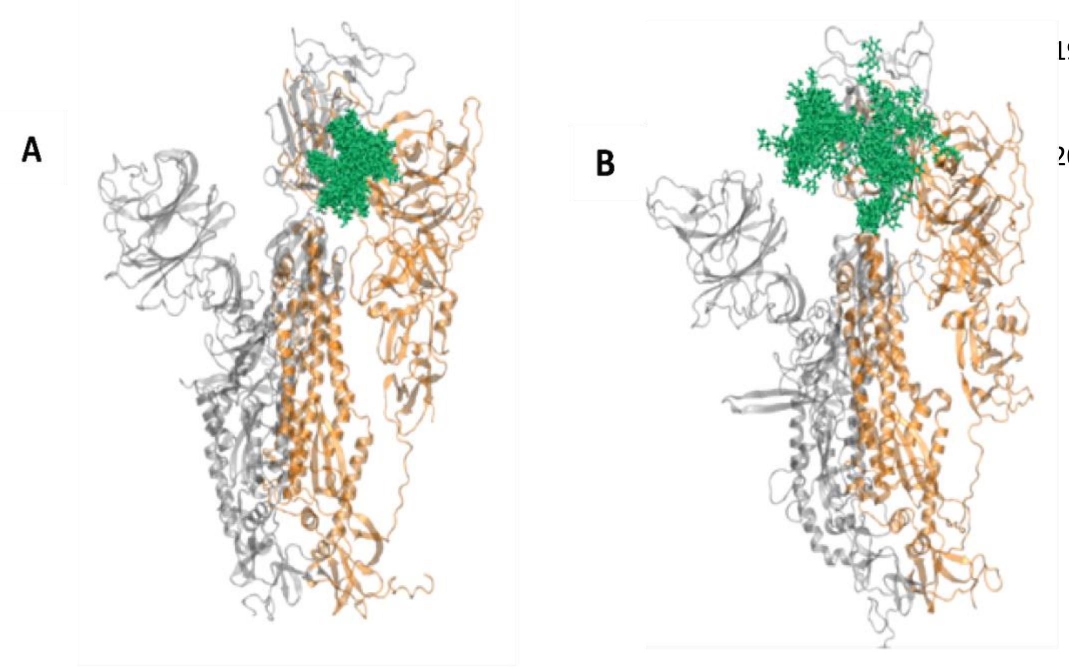


**SI Figure 8.** Representation of glycan N343 present on the RBD for both the **A.** Cleaved and the **B.** Uncleaved conformations
